## Supplementary Materials for "AI-predicted protein deformation encodes energy landscape perturbation"

ononoffoffononoffoff

### Contents

1. PDB Structure Data 1
  A. How many structures are needed to average? 1
  B. pLDDT is less informative of stability than strain. 1
  C. Insufficient PDB data to measure empirical strain-stability correlation. 1
  D. Construction of a reduced ThermoMutDB set 1

2. Linking spatial deformation and energy using elastic network models 1

### 1. PDB Structure Data

#### A. How many structures are needed to average?

Since we found that correlations between $S_n$ and $\Delta\Delta G$ are higher when using averaged AF-predicted structures, a pertinent practical question is, how many structures should one use? We looked at the correlations (both $\Delta\Delta G$ and $|\Delta\Delta G|$) as a function of the number of structures used to create average structures, finding that most of the improvement is seen when using only five structures instead of one (Fig. 11). We suggest that using 20 structures is sufficient to achieve good performance.

#### B. pLDDT is less informative of stability than strain.

Since AF's predicted confidence measure, pLDDT, is a good indicator of disorder, we also investigated whether changes in pLDDT are informative of changes in stability. We compared changes in pLDDT at the mutated site $m$, changes in the average pLDDT (across the entire protein), for both single and averaged structures, against both $\Delta\Delta G$ and $|\Delta\Delta G|$. We find poor correlations between changes in pLDDT and $\Delta\Delta G$ (Fig. 10), in agreement with Pak *et al.* (2023).

#### C. Insufficient PDB data to measure empirical strain-stability correlation.

We sought to repeat our analysis using experimental structures from the Protein Data Bank (PDB). We only found $\sim 100$ entries which have both wild-type (WT) and mutant experimental structures. We removed any NMR structures, and only considered pairs with the same oligomeric state and same bound substrates. This left us with 36 pairs of PDB structures, which is too few to study statistical correlations. It is possible that there are sufficient $\Delta\Delta G$ measurements across other data sets, but the limit of unique mutations is determined by the number of possible structure pairs that differ by a single amino acid substitution. In a previous study (McBride *et al.*, 2023) we only found $\sim$1,000 of these, so there is some hope that this analysis can be performed if the other datasets have enough $\Delta\Delta G$ measurements.

#### D. Construction of a reduced ThermoMutDB set

Our original approach was to use the ThermoMutDB subset provided by Pak *et al.* (2023), however we decided to work exclusively with the original ThermoMutDB data, for two of reasons. We found that the "TMDB_ID" values do not always match with the original ThermoMutDB ID numbers. We also found a case where a mutant (P00644, L89V) was assigned an incorrect mutation position; in this case, both the PDB SEQRES and Uniprot sequences have a lysine at position 89, but the correct mutation code should be L171V.

We found several issues in the ThermoMutDB that led to exclusion or correction of entries. We found that for some entries, the reference protein for which the mutation was defined (and $\Delta\Delta G$ was calculated) was not the WT sequence found in Uniprot; (Consortium, 2018; Isom *et al.*, 2008, 2010; Merkel *et al.*, 1999) we excluded these. We found duplications of entries, where measurements from one paper were referenced in several other papers, yet both were included; (Robinson *et al.*, 2018) we excluded these entries. We found that one measurement (Y27A) in one source, (Shortle *et al.*, 1990) and all measurements in another source (Carra and Privalov, 1995) were included with the incorrect sign; we corrected these.

### 2. Linking spatial deformation and energy using elastic network models

It is instructive to examine the energy landscape within a framework of effective elasticity theory, which allows us to link the scale of free energy changes with the underlying structural changes. The generic relation for a single spring is $U = \frac{1}{2}K\Delta r^2$, where $K$ is a spring constant and $\Delta r$ is the deviation from the spring's equilibrium length. A generalization to 3D continuum material is a quadratic form for energy density, $u = \frac{1}{2}\mathbf{k}_{\alpha\beta\gamma\delta}\mathbf{S}_{\alpha\beta}\mathbf{S}_{\gamma\delta}$, where $\mathbf{S}$ is the $3\times3$ strain tensor, $\mathbf{k}$ is the $3\times3\times3\times3$ elasticity tensor (Landau *et al.*, 1986), and we use Einstein's summation convention.

To obtain a similar effective strain-energy relation for the protein, we model a protein as an elastic network where amino acids are connected by homogeneous springs with elastic constant $K$ between their $C_\alpha$ positions. Two $C_\alpha$ positions $i$ and $j$ are connected if they are within $\gamma$ Å from each other, $|\mathbf{r}_{ij}| < \gamma$. To measure the deformation, we use our previously-developed metric called the local distance difference (LDD), (McBride *et al.*, 2023) which is the L2-norm of the vector of changes in the $C_\alpha - C_\alpha$ distances between neighbors,

$$\mathrm{LDD}_i = \sqrt{\sum_{j=1}^{n_i} \left(|\mathbf{r}_{ij}| - |\mathbf{r}'_{ij}|\right)^2}\,.$$

Here, $\mathbf{r}_{ij}$ and $\mathbf{r}'_{ij}$ are the vectors between residue $i$ and neighbor $j$ in, respectively, a reference structure and a target structure, and the summation is over the $n_i$ neighbors $j$ for which $|\mathbf{r}_{ij}| < \gamma$. In practice, we average $|r_{ij}|$ and $|r'_{ij}|$ over many structures to get an estimate of the equilibrium distance between neighboring residues. LDD is highly correlated with effective strain (McBride *et al.*, 2023), and thus is also correlated with $\Delta\Delta G$ (Table I). Then, the overall elastic energy of the network is simply the sum

$$U = \frac{1}{4} K \sum_{i=1}^{N} \sum_{j=1}^{n_i} \left(|\mathbf{r}_{ij}| - |\mathbf{r}'_{ij}|\right)^2 = \frac{1}{4} K \sum_{i=1}^{N} \mathrm{LDD}_i^2\,, \qquad (1)$$

where the additional $\frac{1}{2}$ factor accounts for counting twice each $i - j$ bond.

To get a rough estimate for the typical values of *effective* spring constant $K$ that describes the energy landscape of the protein, we first compute the LDD by comparing wild-type and mutated AF-predicted structures, exactly in the manner in which the effective strain is computed (see App. B in manuscript). Next, we equate the change in stability $\Delta\Delta G$ to the elastic energy $U$ in Eq. (1), from which we obtain an approximation for $K$,

$$K = \frac{4\,\Delta\Delta G}{\sum_i \mathrm{LDD}_i^2}\,. \qquad (2)$$

It is essential to note that $\Delta\Delta G$ includes significant contributions from changes in the entropy and the topology of the bond network (bonds may break and reform), which may sometimes decrease the overall energy. Moreover, many bonds in the equilibrium conformation of the protein are “frustrated” (Ferreiro *et al.*, 2014), *i.e.* shorter or longer than their equilibrium length. Hence, deformation can release some of the elastic energy in these pre-stressed bonds. For all these reasons, the estimated spring constant $K$ from Eq.(2) is only an effective parameter, which serves as a ballpark figure for the actual strength of elastic interactions between amino acids.

Using this method, we estimate $K$ for each of the 46 protein families for which we have 10 or more $\Delta\Delta G$ values. We calculate the sum of squared $\mathrm{LDD}_i$ for each protein in a family, fit a straight line between this and $\Delta\Delta G$ using Eq.(1) with an intercept at zero, and get $K$ from the gradient. As expected, the value of $K$ depends strongly on the topology of the assumed bond network, in which the number of springs is determined by the neighborhood radius $\gamma$. Choosing $\gamma \sim 6-7$ Å corresponds to bonds between amino acids that are likely to be directly in physical contact. At this range, we find an average spring strength of $K \sim 10\,\mathrm{kcal/mol/Å}^2$ (Fig. 9). Choosing Larger neighborhood radii $\gamma \sim 10-11$ Å, as commonly assumed in elastic network models (McBride *et al.*, 2022), yields lower average values of $K \sim 1\,\mathrm{kcal/mol/Å}^2$. The effective $K$ values vary broadly, with a standard error of about one order of magnitude, reflecting the diverse, intricate geometry of protein energy landscapes. Even in a single protein family, the highly anisotropic landscape exhibits a wide distribution of spring constants. Certain directions in the space of conformations are very flat as the corresponding deformation hardly changes $\Delta\Delta G$, while other directions are very steep with large $\Delta\Delta G$ changes (McBride *et al.*, 2022).

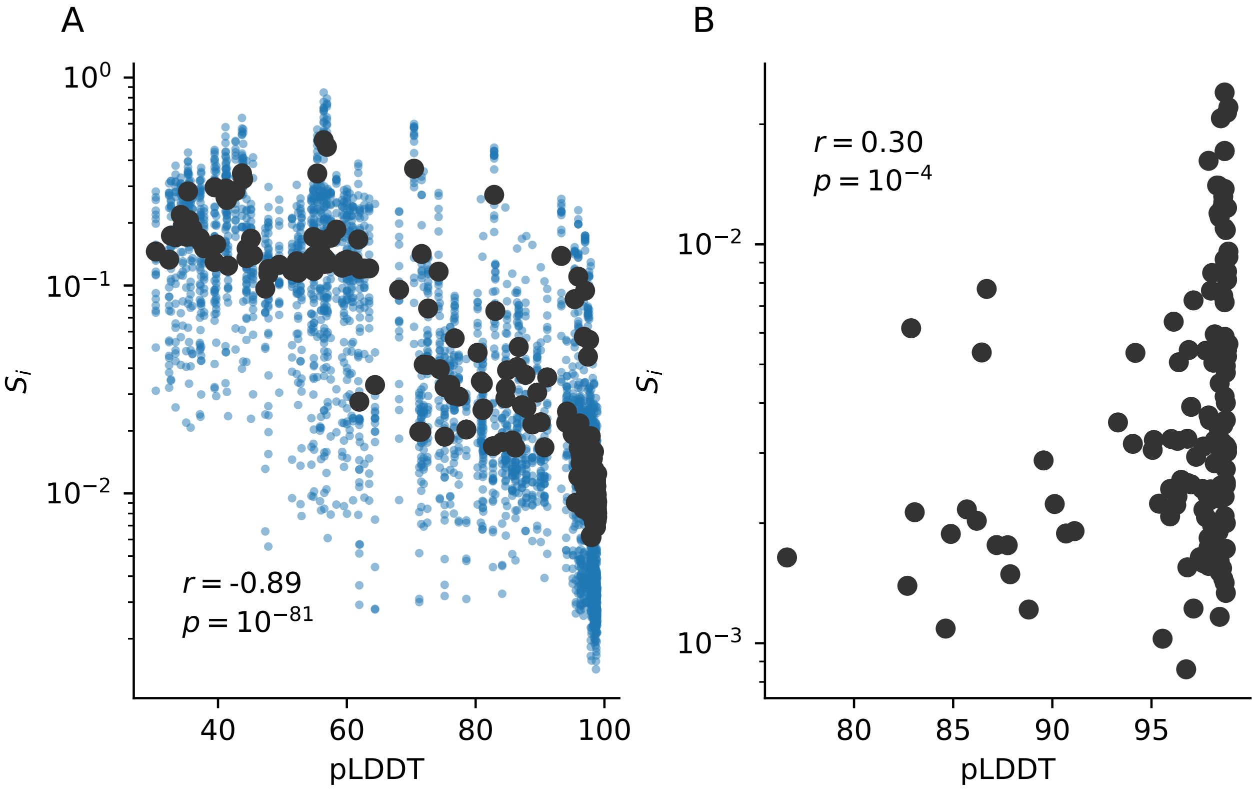


FIG. 1 **AF predictions are more variable in regions with low pLDDT.** Strain per residue $S_i$ vs. pLDDT per residue for: (A) Strain calculated between repeat AF predictions of the same protein sequence; blue dots are shown for different pairs of single structures; black circles indicate means across multiple pairs. (B) Strain calculated for pairs of averaged structures, whose sequences differ by a single mutation; residues with pLDDT $< 70$ are excluded. Pearson's correlation coefficient and p value are shown.

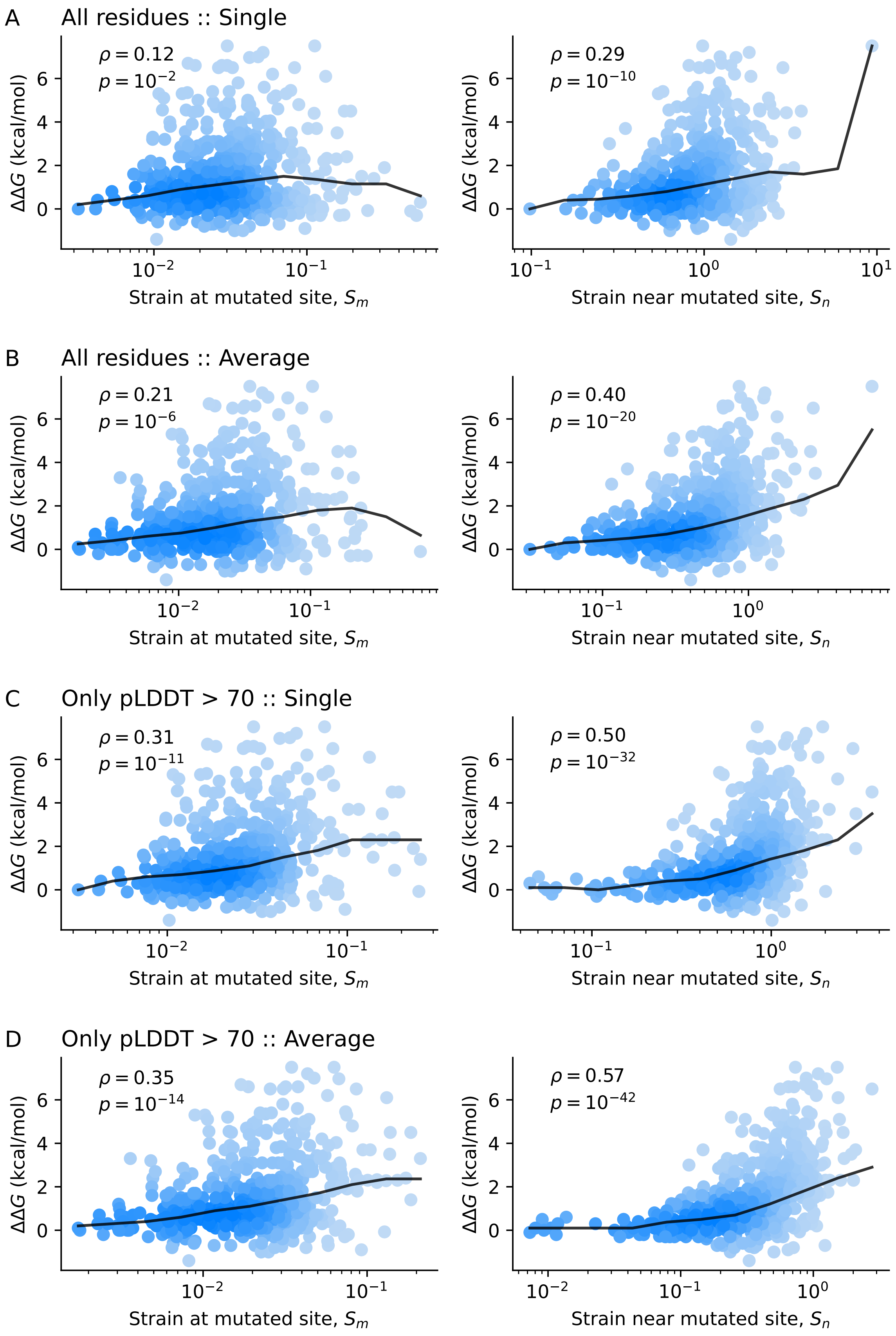


FIG. 2 **Improving precision of mutation effect prediction.** Strain at the mutated site $m$, $S_m$ (left), and near the mutated site, $S_n$ (right), for four different methods of calculating strain: (A) All residues and pairs of single structures. (B) All residues and pairs of averaged structures. (C) Only residues with pLDDT $>$ 70 and pairs of single structures. (D) Only residues with pLDDT $>$ 70 and pairs of averaged structures. Spearman's $\rho$ and p value are shown.

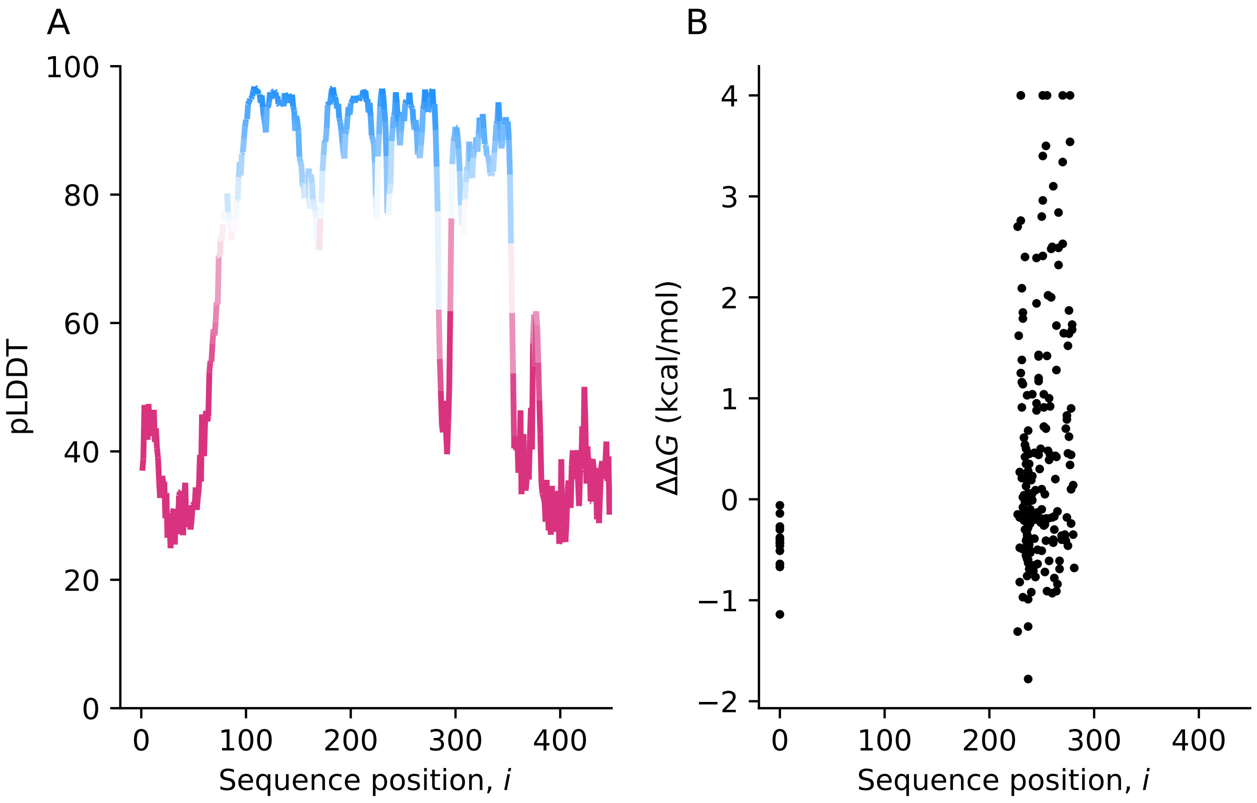


FIG. 3 **Biased sampling of IgG-binding protein G (IgGb).** A: pLDDT vs sequence position $i$ for IgGb. B: Distribution of mutations in the ThermoMutDB for IgGb along the sequence, and the corresponding $\Delta\Delta G$ values.

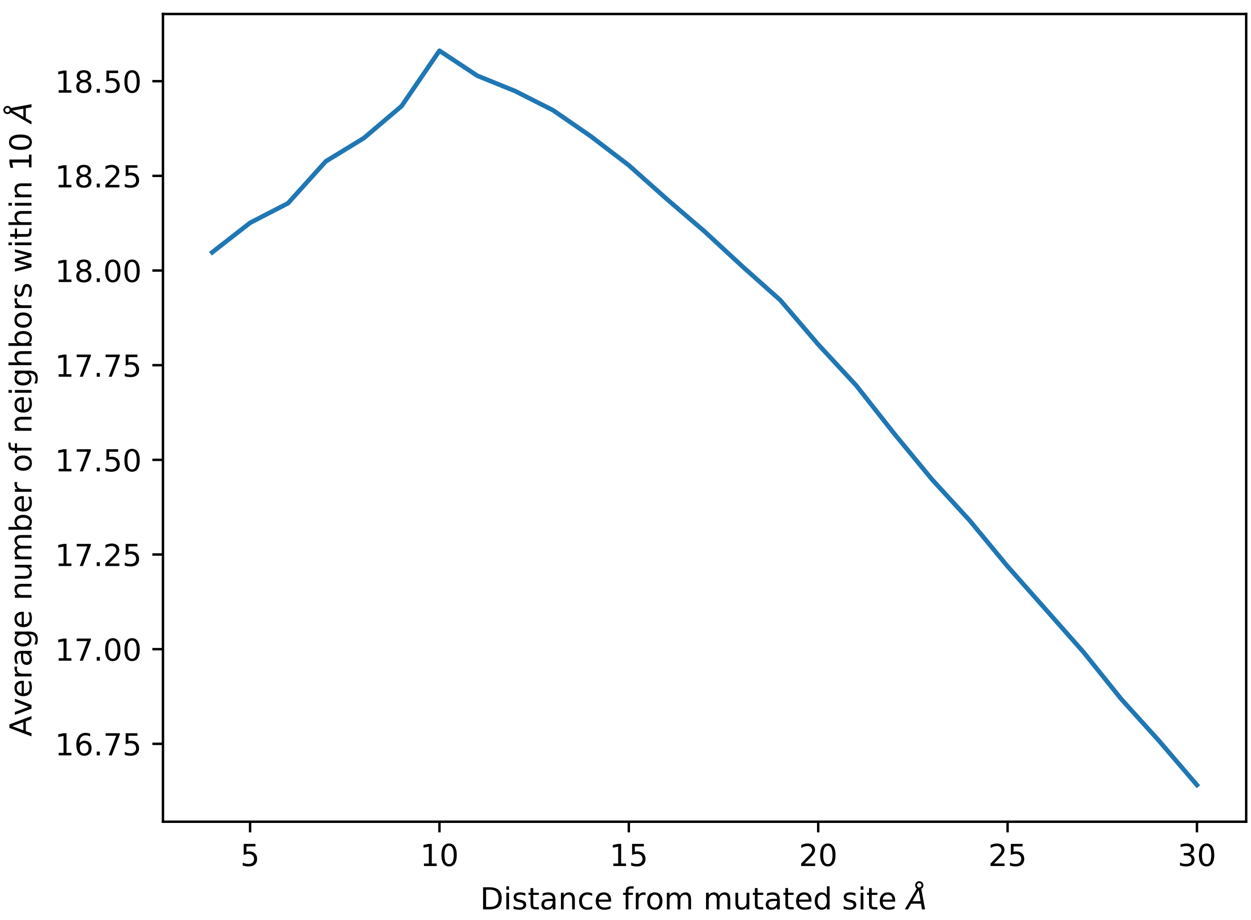


FIG. 4 **The most buried residues tend to be within 15 Å of other residues.** For each protein we measure the number of neighbors per residue. Neighbors are defined as the set of residues whose $C_\alpha$ distances are within 10 Å. Plot shows average number of neighbors of residues as a function of the distance to the mutated site. Average is calculated for all unique proteins in our reduced ThermoMutDB sample.

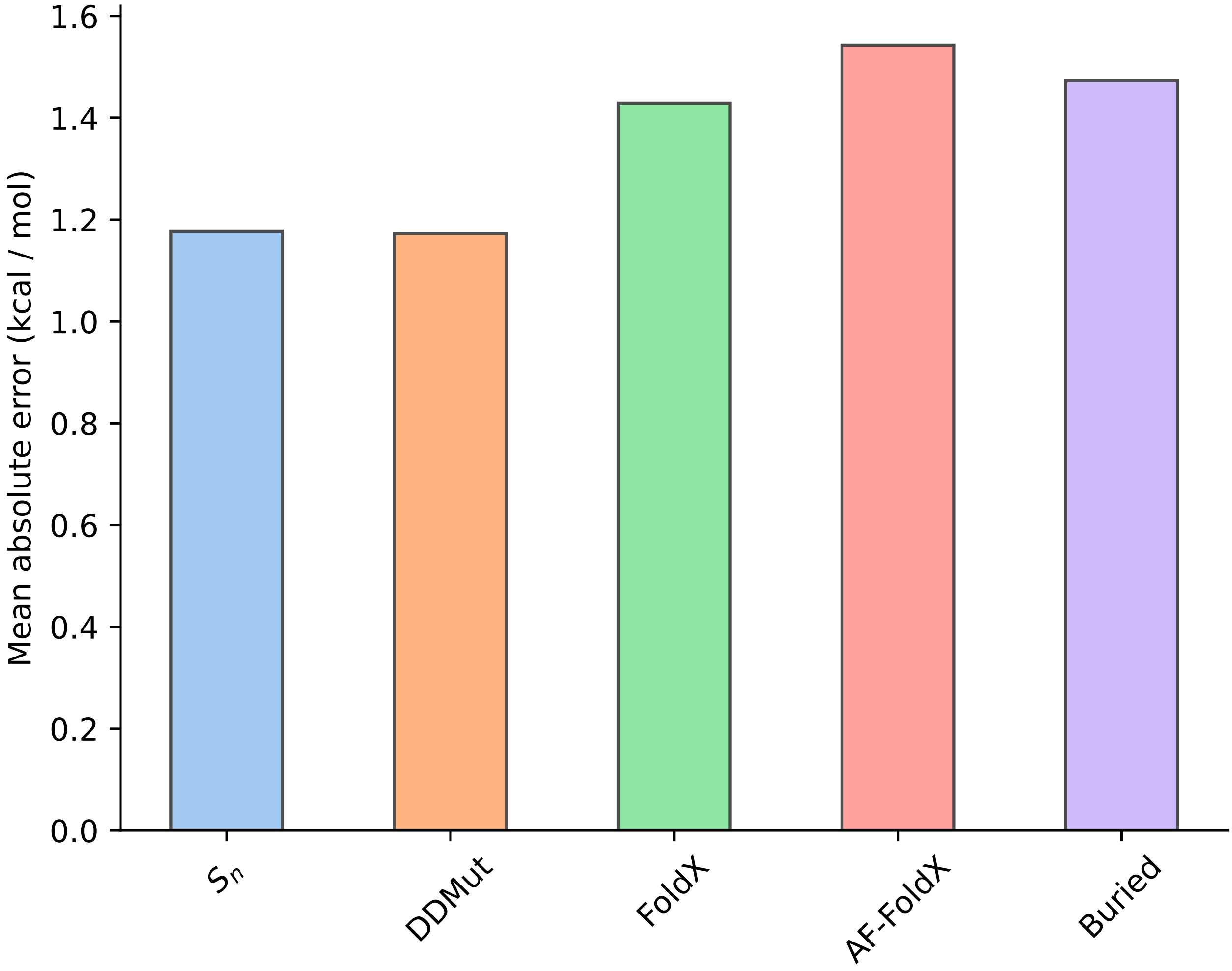


FIG. 5 Mean absolute error of linear fits to $\Delta\Delta G$ using various methods.

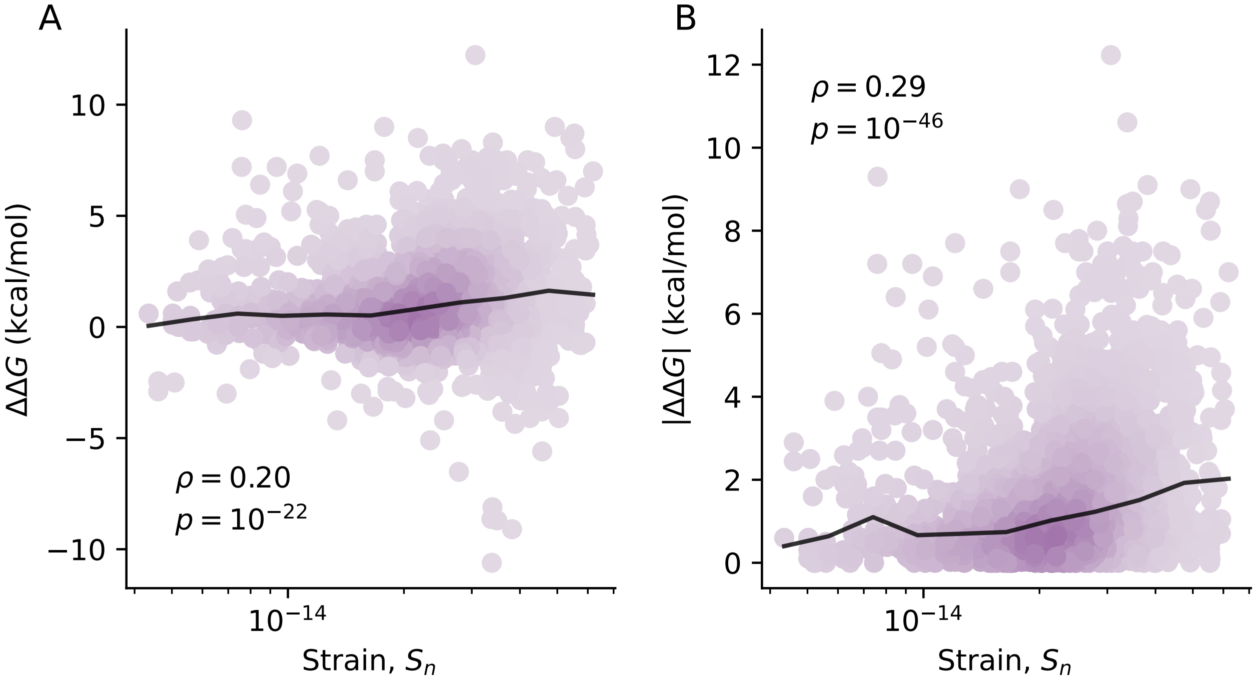


FIG. 6 **$S_n$–$\Delta\Delta G$ correlations calculated using FoldX structures.** Strain near mutated site $S_n$ vs. $\Delta\Delta G$ (A) and magnitude of stability change $|\Delta\Delta G|$ for all mutations in our reduced ThermoMutDB sample of 2,499 unique mutants. Spearman's $\rho$ and p value are shown.

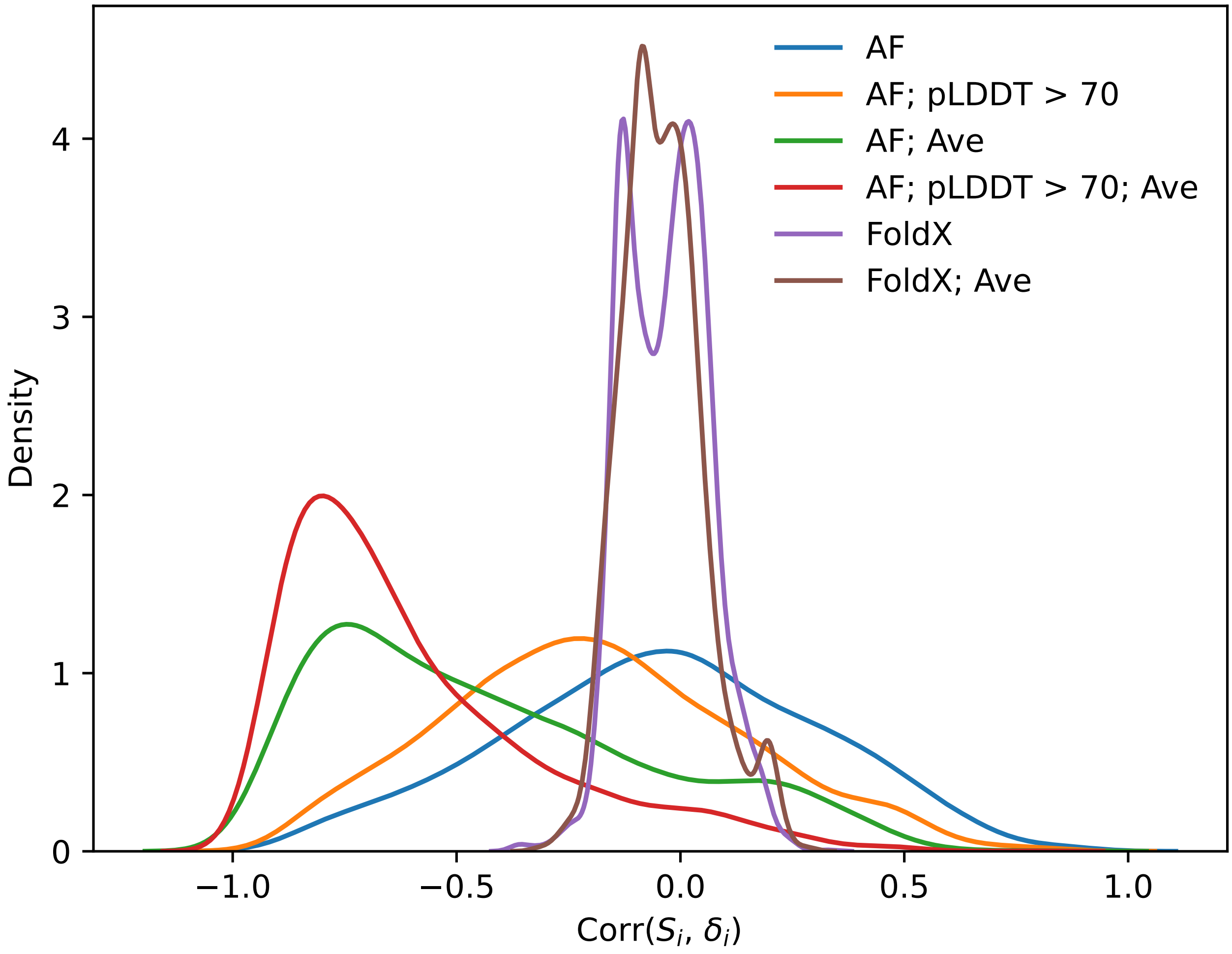


FIG. 7 **Correlation between distance from mutated site $\delta_i$ and $S_i$** for all proteins in our reduced ThermoMutDB sample. Different distributions are shown for shear calculated using: AF-predicted structures, all residues and pairs of single structures (AF). AF-predicted structures, only residues with pLDDT $>$ 70 and pairs of single structures (AF; pLDDT $>$ 70). AF-predicted structures, all residues and pairs of averaged structures (AF; Ave). AF-predicted structures, only residues with pLDDT $>$ 70 and pairs of averaged structures (AF; pLDDT $>$ 70; Ave). FoldX-predicted structures, all residues and pairs of single structures (FoldX). FoldX-predicted structures, all residues and pairs of averaged structures (FoldX; Ave).

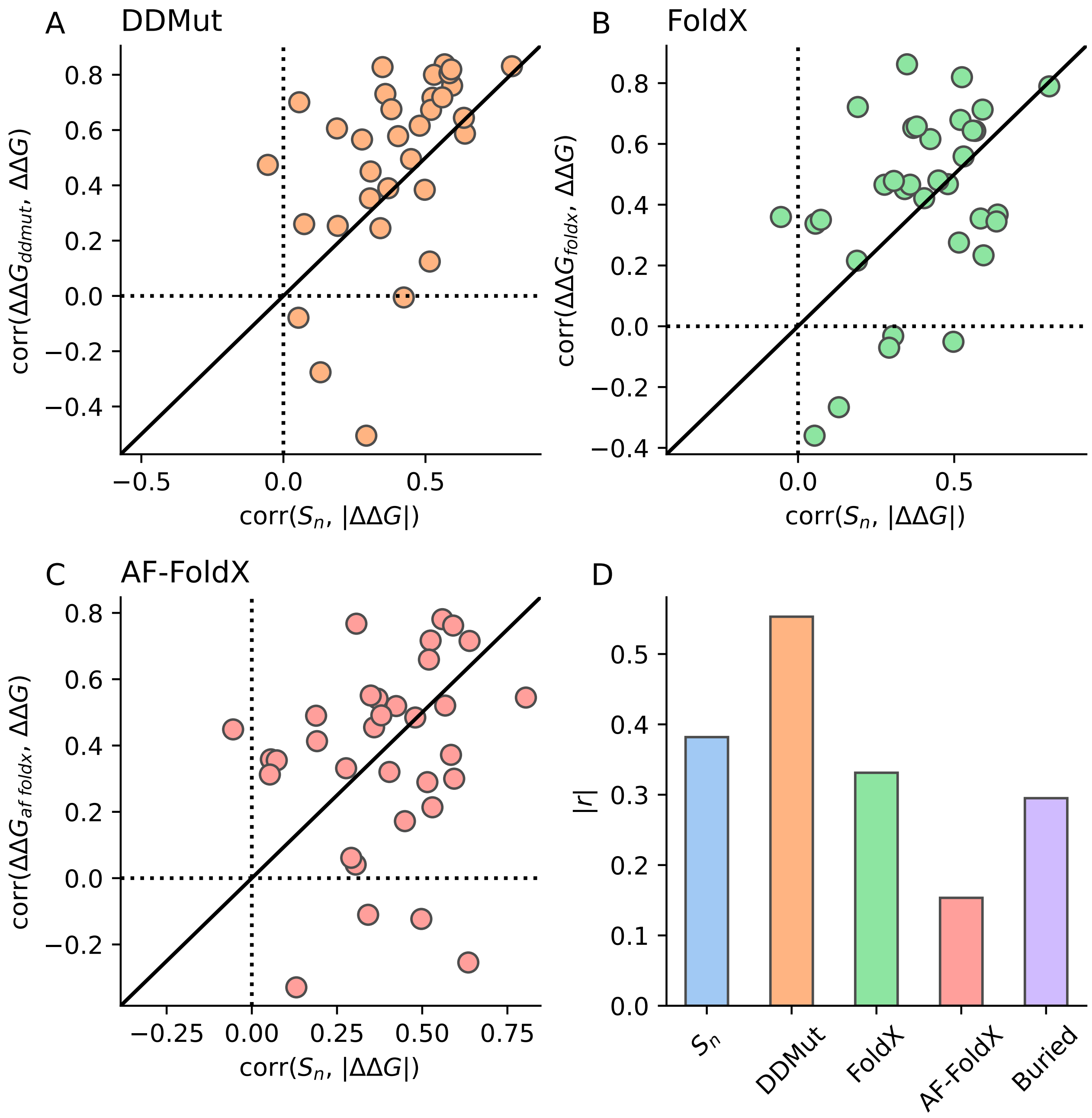


FIG. 8 **Comparison with state-of-the-art algorithms.** A-C: Correlation (Pearson's $r$) between strain $S_n$ and magnitude of stability change $|\Delta\Delta G|$, vs. correlation between $\Delta\Delta G$ and predictions of DDMut (A), FoldX (B), and FoldX using AF-predicted structures (C); separate points are shown for each of the 40 most-common proteins. D: Correlations (as above) for the full sample of 2,499 unique mutants. Correlation between number of neighbors within 10 Å of the mutated site is shown as "Buried".

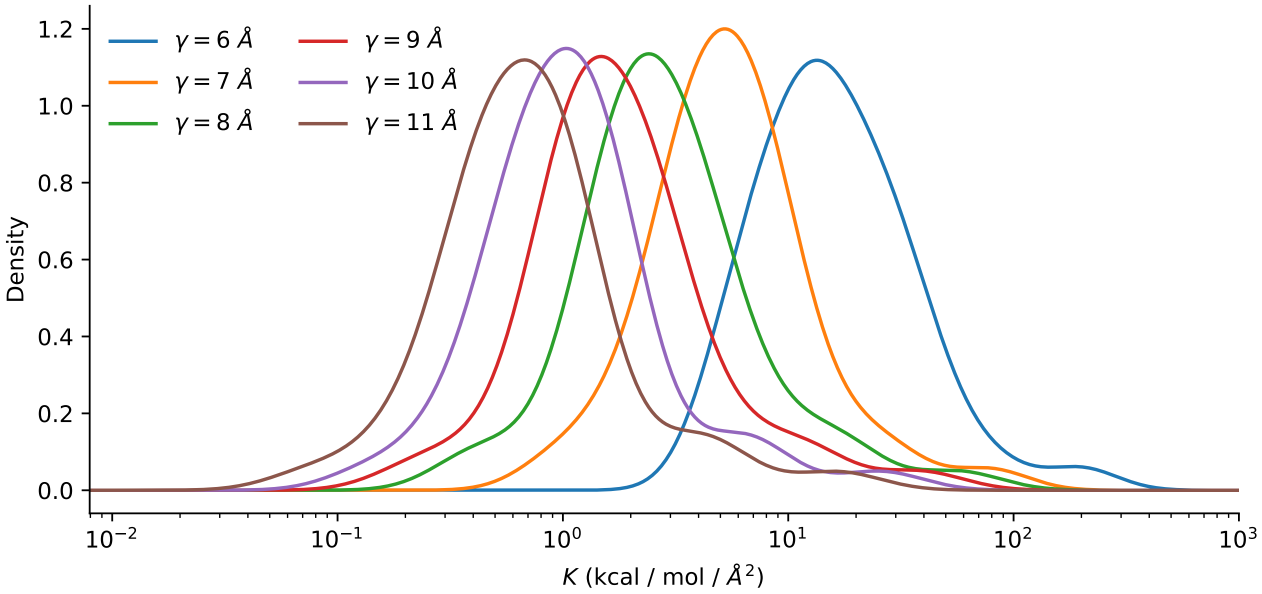


FIG. 9 **Computation of effective spring constant from stability changes.** Distribution of homogeneous spring constant $K$ inferred from empirical $\Delta\Delta G$ and predicted deformation using an elastic network model (Eq.(2)). Different values of $K$ are obtained for different elastic network topologies, defined by the neighborhood cutoff radius $\gamma$ Å.

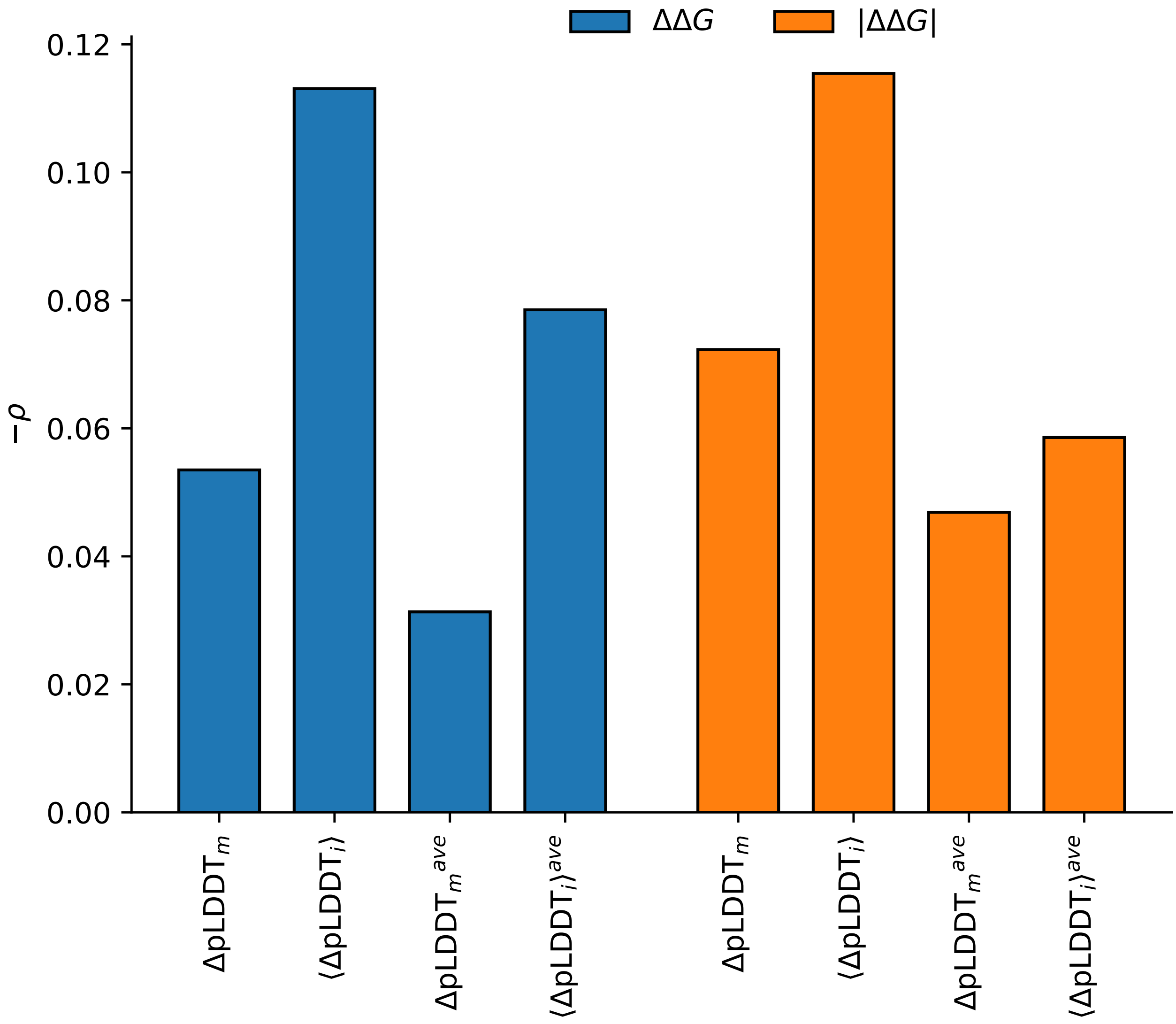


FIG. 10 **ΔpLDDT is a poor predictor of ΔΔ*G*.** Spearman's correlation between differences in pLDDT and stability. Differences in pLDDT are measured in four ways: difference in pLDDT at the mutated site $\Delta\mathrm{pLDDT}_m$; average difference in pLDDT across the whole protein $\langle\Delta\mathrm{pLDDT}_i\rangle$; difference in pLDDT at the mutated site, averaged over 50 pLDDT predictions $\Delta\mathrm{pLDDT}_m^{ave}$; average difference in pLDDT across the whole protein, averaged over 50 pLDDT predictions $\langle\Delta\mathrm{pLDDT}_i\rangle^{ave}$;

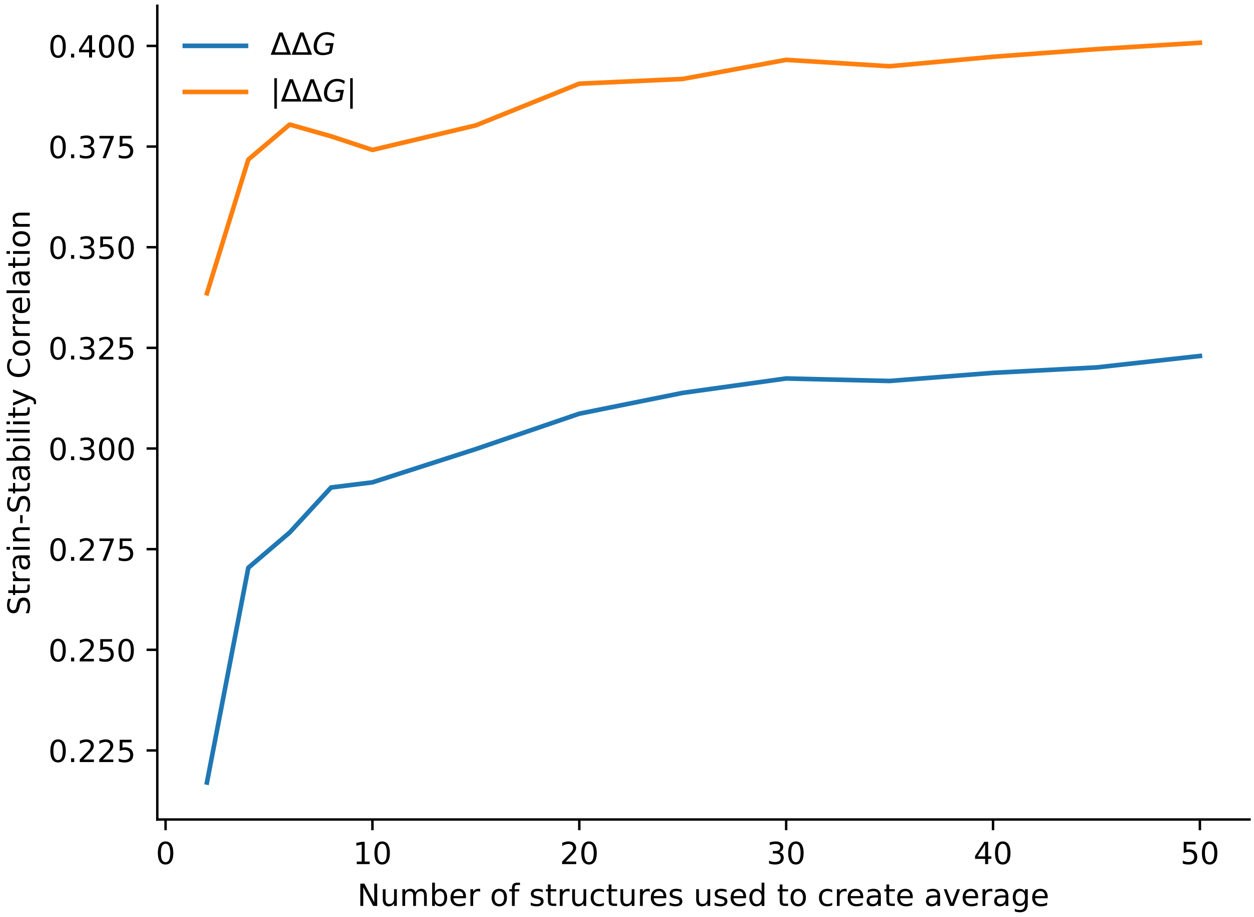


FIG. 11 **Effect of the number of structures used to create averages.** Strain-stability correlation for all mutations in our reduced ThermoMutDB sample of 2,499 unique mutants, as a function of the number of structures used to create average structures.

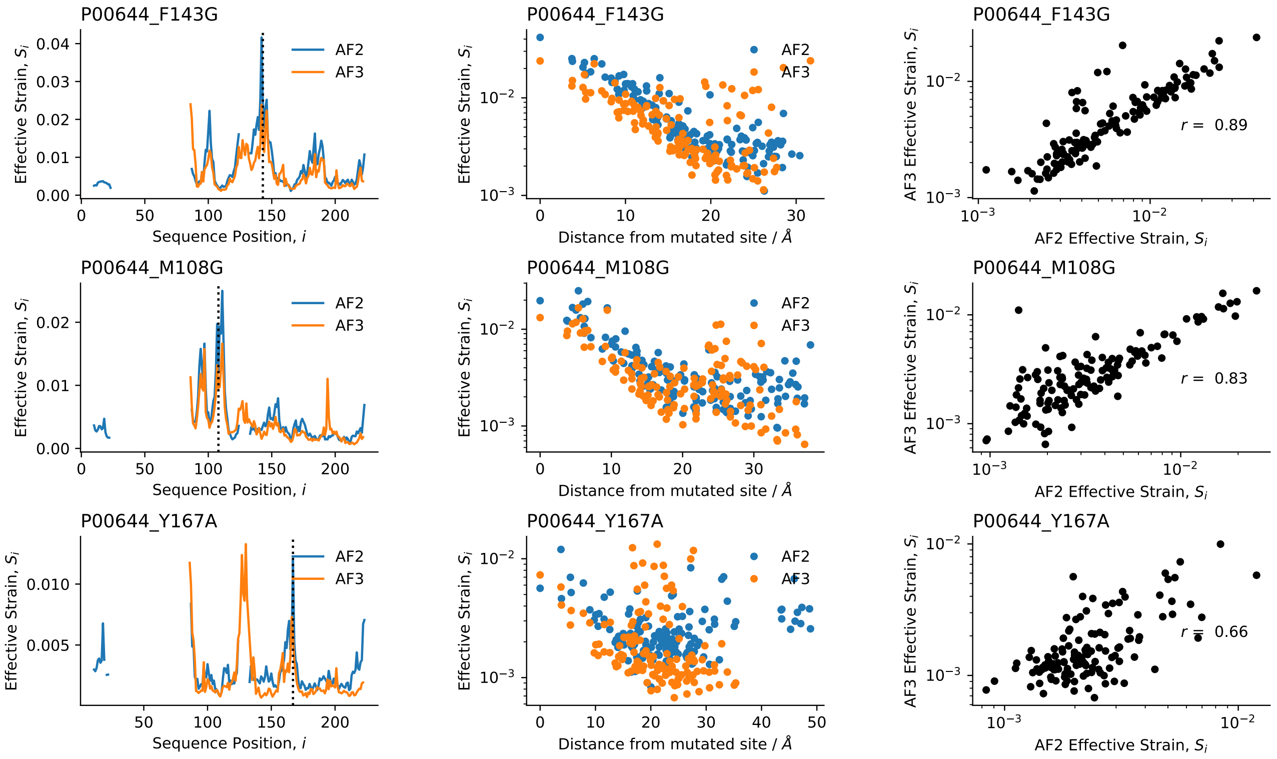


FIG. 12 **Comparison of AlphaFold2 (AF2) and AlphaFold3 (AF3) strain predictions.** Strain is calculated by AF2 and AF3 (Abramson *et al.*, 2024) between Themonuclease WT (P00644) and three mutants: F143G (top), M108G (middle) and Y167A (bottom). Left: Strain (AF2 and AF3) as a function of sequence position. Center: distance from the mutated site (center). Right: AF3-predicted strain against AF2-predicted strain (Pearson's $r$ is shown). The strain values are generally similar, except for some flexible regions where AF2 and AF3 give different pLDDT values (*e.g.*, the loop around residue 128, see SI Fig. 13).

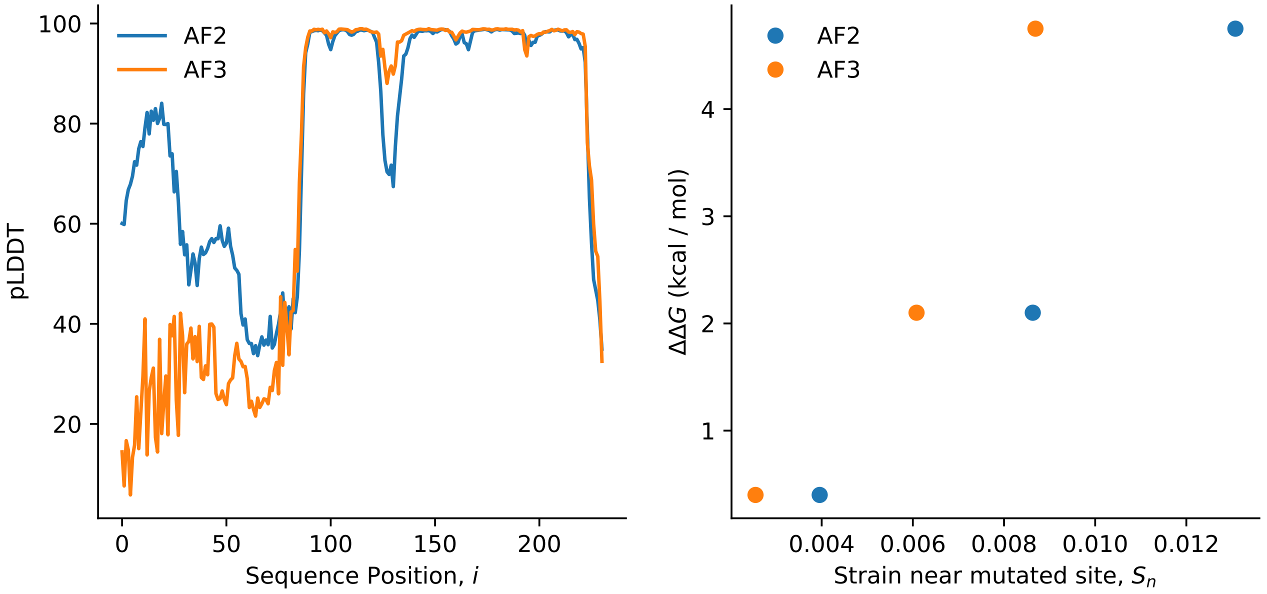


FIG. 13 **Comparison of AF2 and AF3 pLDDT and mutation effect on stability.** Left: pLDDT (AF2 and AF3) vs. sequence position.

TABLE I  Comparison of linear and rank-order correlations between $\Delta\Delta G$ and either Effective Strain or LDD (summation over residues near the mutated site), for main text Figures 1-3.

| Figure | Correlation | Strain | LDD |
|---|---|---|---|
| Fig1F | Linear | 0.35 | 0.4 |
| Fig1F | Rank Order | 0.35 | 0.41 |
| Fig1G | Linear | 0.52 | 0.51 |
| Fig1G | Rank Order | 0.57 | 0.55 |
| Fig2A (i) | Linear | 0.44 | 0.42 |
| Fig2A (i) | Rank Order | 0.37 | 0.36 |
| Fig2A (ii) | Linear | 0.6 | 0.59 |
| Fig2A (ii) | Rank Order | 0.61 | 0.59 |
| Fig2A (iii) | Linear | 0.48 | 0.45 |
| Fig2A (iii) | Rank Order | 0.52 | 0.47 |
| Fig2A (iv) | Linear | 0.5 | 0.49 |
| Fig2A (iv) | Rank Order | 0.57 | 0.54 |
| Fig3A | Linear | 0.31 | 0.29 |
| Fig3A | Rank Order | 0.36 | 0.34 |
| Fig3B | Linear | 0.37 | 0.35 |
| Fig3B | Rank Order | 0.41 | 0.4 |
